# Distance-stratified benchmarking of variant-effect predictors on *β*-lactamase deep mutational scanning data

**DOI:** 10.64898/2026.11.24.768893

**Authors:** Akshaya Dhakal, Subash Acharya, Gungun Chand

## Abstract

Mutations far from the active site of antibiotic-resistance enzymes can change resistance, but it is unclear what information predicts their effects. We asked how the performance of a protein language model (ESM2), a structure-based stability predictor (ThermoMPNN) and simple structural descriptors changes with distance from the catalytic residues, using deep mutational scanning data for the *β*-lactamases TEM-1 (four assays) and VIM-2 (one assay). We measured Spearman correlation and AUROC within distance bins, with bootstrap intervals over positions. In TEM-1, ESM2 accuracy fell with distance (Stiffler assay: *ρ* = 0.72 below 8 Å and 0.42 beyond 20 Å), while the stability score rose (*ρ* = 0.41 to 0.70). Adding stability to a model of ESM2, solvent accessibility and neighbour count improved cross-validated *ρ* by 0.02 below 8 Å and 0.10 beyond 20 Å, and the combined model did not decline with distance. The pattern held across binning schemes, in a continuous-distance model, and when any one of five contiguous distal stretches was removed. Two TEM-1 assays are nearly duplicates, and the Deng assay was too noisy to be informative. VIM-2 has only seven positions beyond 20 Å and showed no detectable trend within 20 Å, so it neither supports nor contradicts the TEM-1 pattern. Nine distal TEM-1 positions remained more sensitive than the models predicted. In TEM-1, predicted stability was more informative than the language model far from the catalytic site, and the language model was stronger near it. Distance-stratified reporting could add to variant-effect benchmarks. The study is exploratory and uses one structure and one phenotype per assay.

## 1 Introduction

*β*-Lactamases are among the most clinically important antibiotic-resistance enzymes (1). TEM-1, a class A serine *β*-lactamase, hydrolyses penicillin-type antibiotics through an acyl-enzyme intermediate formed at Ser70, with Lys73, Ser130 and Glu166 contributing to catalysis (residue numbers in Ambler numbering) (2). VIM-2 is a class B1 metallo-*β*-lactamase that uses a dinuclear zinc centre instead of a catalytic serine (3). Deep mutational scanning (DMS) (4) has measured the effect of thousands of single amino-acid substitutions on the resistance phenotype of both enzymes: four independent experiments on TEM-1 (5–8) and one on VIM-2 (9). These datasets are curated in ProteinGym, a benchmark for variant-effect predictors (10). A basic question about such data is how far from the catalytic site a mutation can be and still change the resistance phenotype, and what kind of information is needed to predict it. Mutations at or near the active site are expected to act directly on catalysis or substrate binding. Mutations far from the active site could act in at least two ways. They could change the amount of properly folded enzyme, through effects on folding stability, expression or export (11). Or they could act more specifically, for example by altering protein dynamics or long-range communication to the active site (12, 13). These explanations predict different things about which computational signals should work. Growth-based DMS readouts combine catalytic activity with stability, expression and translocation, so the measured quantity is a resistance phenotype and not a catalytic rate constant (14). We use that term throughout. Two families of predictors address different parts of this problem. Protein language models and conservation-based methods capture evolutionary constraint, which should include catalytic and functional requirements (15–17). Structure-based predictors of folding stability capture biophysical effects on the folded state, and recent models are trained on large-scale stability measurements (18, 19). Benchmarks such as ProteinGym usually report one correlation per protein, which averages over all positions regardless of their distance from the active site. A single average cannot show where along the structure a predictor succeeds or fails, or whether stability information matters more far from the active site. Related work has addressed parts of this question. Benchmarks against DMS data have compared predictors across many proteins (20), and predicted stability effects correlate best with abundance-based DMS phenotypes (21). In enzymes, the predictability of variant effects by supervised models depends on structural features, including proximity to the active site (22). Here we ask how the performance of a language model (ESM2), a stability predictor (ThermoMPNN) and simple structural descriptors changes with distance from the catalytic residues in *β*-lactamase DMS data. Our hypothesis, stated before analysis, was that language-model performance would decline with distance and that predicted stability would explain a growing share of the remaining variation. The alternative was that performance would not depend on distance, or that stability predictions would add no information beyond the language model and burial. We tested this with four independent TEM-1 assays, which also provide a replicate-based estimate of experimental agreement, and with VIM-2 as a structurally and mechanistically different enzyme. Confidence intervals come from bootstrap resampling of positions, and we checked the main result against alternative binning schemes, alternative definitions of a deleterious variant, and the removal of individual contiguous stretches of the protein. In TEM-1, language-model accuracy fell with distance while predicted stability added information that grew with distance. In VIM-2, which has few positions beyond 20 Å, we did not detect a distance trend within the range it samples.

## 2 Methods

### 2.1 Dataset

We used single-substitutes scores from ProteinGym v1.3 (10); four TEM-1 assays (Deng 2012 (5), Jacquier 2013 (6), Firnberg 2014 (7) and Stiffler 2015 (8); 4996, 989, 4783 and 4996 variants) and one VIM-2 assay (Chen 2020(9); 5004 variants; meropenem at 0.031 *µ*g*/*mL, 37 ^*°*^C, according to the ProteinGym label). Higher scores mean higher fitness. Deleterious variants were defined by ProteinGym’s binary label (direction checked from the class mean) and, as secondary definitions, as the lowest 20%, 30% or 40% of scores within an assay. The Firnberg and Stiffler assay agree strongly (Spearman *ρ* =0.94) and are not independent; the Deng assay agree less well(*ρ* = 0.57-0.61), so we treat it as noisier.

### 2.2 Structures, numbering and distance

We used PDB 1BTL (23) for TEM-1 and PDB 6YRP (24) for VIM-2 (an inhibitor-bound structure at 1.95 Å; the inhibitor was removed), chain A, protein atoms only. We inferred the offset between ProteinGym and sequence numbering from the wild-type residues and aligned each reference sequence to its structure (Biopython PairwiseAligner (25)). For TEM-1, all 263 structure residues mapped (identity 0.992); the two residues that differ from 1BTL (ProteinGym positions 82 and 182; Ambler 84 and 184) were excluded from stability-based analyses. For VIM-2, 231 of 266 positions mapped (identity 0.996); positions 1-31 and 263-266 have no structure (13.2% of variants) and were excluded from distance analyses, and position 185 (a mismatch) from stability-based analyses. TEM-1 residues are given in Ambler numbering (2). The site was Ser70, Lys73, Ser130 and Glu166 for TEM-1 (identities checked in the structure). For VIM-2 we used the two zinc ions that each have three protein ligands within 2.8 Å (His116, His118, His196; Asp120, Cys221, His263) and excluded a third zinc with a single ligand (His285) as a surface site. Distance is the minimum heavy-atom distance between a residue and the site. Relative solvent accessibility (RSA) came from the Shrake-Rupley algorithm (26) on the protein-only structure, normalised by maximum accessibilities (27); neighbours were Cα atoms within 10 Å. An earlier pipeline version computed TEM-1 RSA with crystal-lographic waters present (mean RSA 0.136 instead of 0.231); this was corrected and all RSA-dependent analyses were rerun.

### 2.3 Predictors

The language-model score was ESM2 (650M parameters) (15): the log-probability of the mutant residue minus that of the wild-type residue, with the position masked and no fine-tuning. The stability score came from ThermoMPNN (19), which builds on ProteinMPNN (28), trained on large-scale stability data (18), using the default checkpoint in site-saturation mode on the protein-only structure. It does not model metal ions. Its output positions were mapped to the structure by alignment (identity 1.000). Positive values were destabilising (buried positions had higher mean values than exposed ones, and proline the highest), so we use the negated value as “stability”. The base model was ESM2 + RSA + neighbours and the full model added stability; both are linear regressions on percentile ranks, fit separately within each distance bin.

### 2.4 Stratification and evaluation

TEM-1 bins were < 8, 8-14, 14-20 and > 20 Å from the site. VIM-2 has only 7 positions beyond 20 Å, so for VIM-2 we used < 8, 8-14 and > 14 Å and applied the same bins to TEM-1 for a matched comparison; this choice followed the bin sizes but preceded any predictor results. We also used position quartiles and a continuous model with ESM2 × distance and stability × distance interactions (distance standardised within each enzyme). Metrics were Spearman correlation and AUROC for deleterious versus tolerated variants within each bin. ESM2, stability and RSA were evaluated directly; the base and full models used out-of-fold predictions from five-fold cross-validation grouped by position (29). Confidence intervals (95%) are bootstrap percentile intervals (30) over positions (300-1,000 resamples) and do not refit the models; far-versus-near differences use independent bootstraps of the two bins. No multiple-comparison correction was applied.

### 2.5 Robustness

We repeated the far-versus-near comparison with alternative cut-offs (6/12/18 and 10/16/22 Å), quartiles, deleterious thresholds of 20%, 30% and 40%, and after removing the TEM-1 catalytic residues or the six VIM-zinc ligands. The 40 TEM-1 positions beyond 20 Å fall in five contiguous stretches (Ambler 26-41, 50-59, 226-229, 251-258 and 281-290), so position-level resampling may understate uncertainty; we therefore left out each stretch in turn and used a block bootstrap over stretches (uncross-validated correlations). Agreement between TEM-1 assays within each bin is reported as a rough indicator of reproducibility, not an exact ceiling; VIM-2 has no replicate.

## 3 Results

We analysed 4,958 (Stiffler (8), Deng (5)), 4,745 (Firnberg(7)) and 979 (Jacquier (6)) TEM-1 variants at 261 positions, and 4,324 VIM-2 variants (9) at 230 positions (Table 1). About half of the variants in each assay were deleterious by ProteinGym’s label (0.49–0.50). Firnberg and Stiffler agreed closely in every distance bin (Spearman =0.88–0.94). Agreement between Jacquier and Stiffler fell from 0.90 below 8 Å to 0.72 beyond 20 Å. Deng agreed poorly with the other assays near the catalytic residues (=0.32–0.34 below 8 Å) and moderately elsewhere (0.52–0.68; Figure S1, Table S1). We therefore treat Deng as the noisiest assay.

**Table 1.** Datasets analysed. Variants and positions after structure mapping and wild-type checks, fraction labelled deleterious, structure, and site definition. TEM-1 site: Ser70, Lys73, Ser130 and Glu166 (Ambler). VIM-2 site: two zinc ions and their six ligands.

| Dataset | Variants | Positions | Frac. deleterious | Structure | Site definition | Source |
|---|---|---|---|---|---|---|
| TEM-1 Stiffler 2015 | 4958 | 261 | 0.50 | 1BTL chain A | Ser70, Lys73, Ser130, Glu166 (Ambler) | ProteinGym BLAT_ECOLX_Stiffler_2015 |
| TEM-1 Firnberg 2014 | 4745 | 261 | 0.50 | 1BTL chain A | Ser70, Lys73, Ser130, Glu166 (Ambler) | ProteinGym BLAT_ECOLX_Firnberg_2014 |
| TEM-1 Deng 2012 | 4958 | 261 | 0.50 | 1BTL chain A | Ser70, Lys73, Ser130, Glu166 (Ambler) | ProteinGym BLAT_ECOLX_Deng_2012 |
| TEM-1 Jacquier 2013 | 979 | 261 | 0.49 | 1BTL chain A | Ser70, Lys73, Ser130, Glu166 (Ambler) | ProteinGym BLAT_ECOLX_Jacquier_2013 |
| VIM-2 Chen 2020 | 4324 | 230 | 0.49 | 6YRP chain A | Zn1: His116, His118, His196; Zn2: Asp120, Cys221, His263 | ProteinGym A4GRB6_PSEAI_Chen_2020 (meropenem 0.031 µg/mL, 37 °C) |

In the Stiffler assay, the correlation between ESM2 (15) and fitness was 0.72 below 8 Å, 0.77 at 8–14 Å, 0.69 at 14–20 Å and 0.42 (95% CI 0.19–0.62) beyond 20 Å (Figure 1, Table 2). Its AUROC for deleterious versus tolerated variants fell from 0.91 to 0.72 (0.57–0.84; Table 2). The Firnberg and Jacquier assays showed the same decline (beyond 20 Å = 0.40 and 0.32; AUROC 0.70 and 0.68; Table 2, Table S6). In the Deng assay ESM2 reached only 0.39 below 8 Å and 0.22 beyond 20 Å (Table S6). With three matched bins, the farminus-near difference in was −0.15 (−0.26 to −0.02) in Stiffler and −0.16 (−0.28 to −0.04) in Firnberg (Figure 2, Table S3). With position quartiles it was −0.19 (−0.35 to −0.05) and −0.22 (−0.38 to −0.07; Table S3).

**Table 2.**
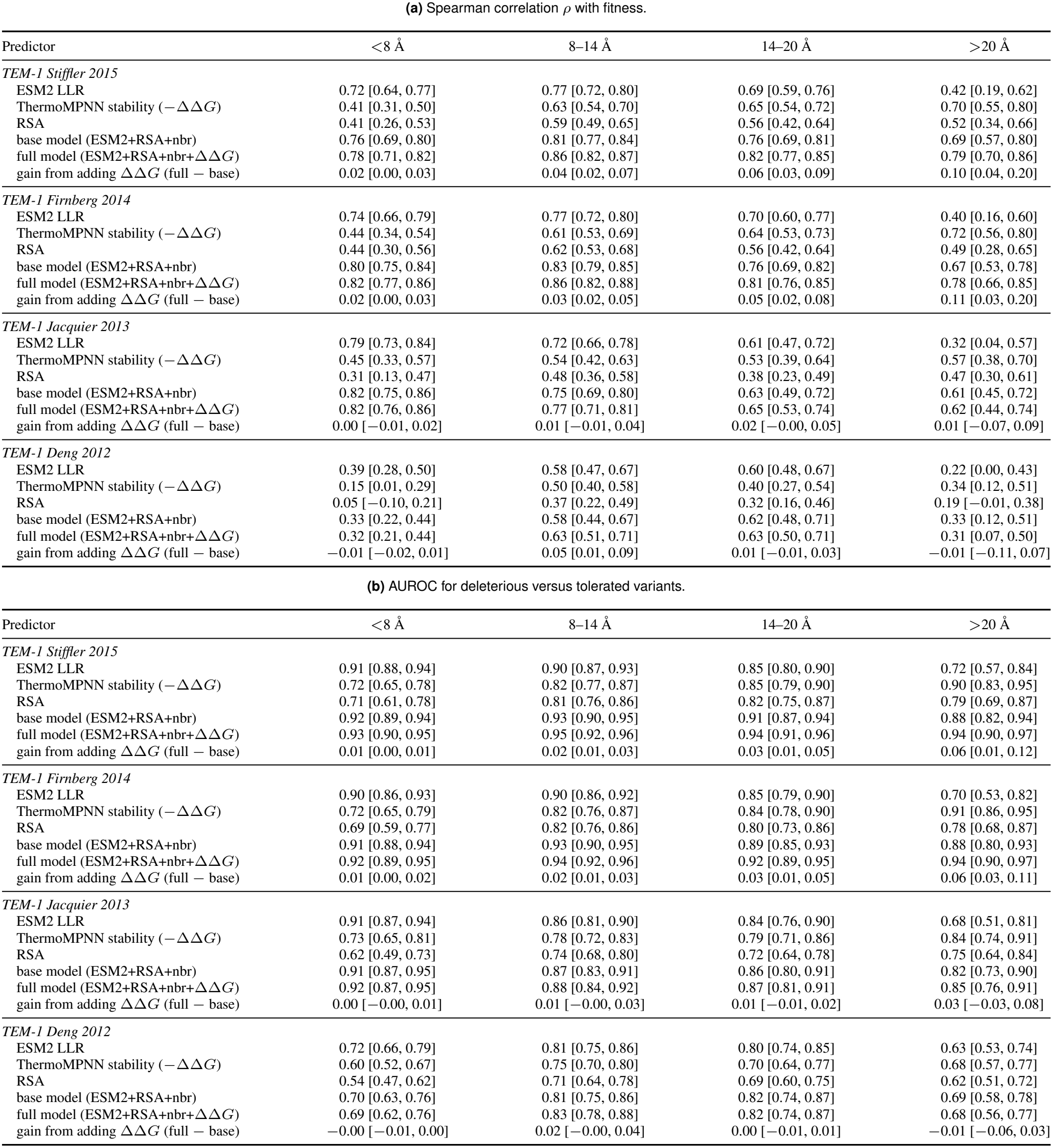
Performance of each predictor by distance to the catalytic residues, for the four TEM-1 assays. Values are point estimates with 95% confidence intervals in brackets. Predictors: ESM2 log-likelihood ratio (LLR), ThermoMPNN stability score, relative solvent accessibility (RSA), a base model (ESM2 + RSA + neighbours, nbr) and a full model (base + stability ΔΔ*G*).

| Predictor | <8 Å | 8–14 Å | 14–20 Å | >20 Å |
|---|---|---|---|---|
| *TEM-1 Stiffler 2015* |  |  |  |  |
| ESM2 LLR | 0.72 [0.64, 0.77] | 0.77 [0.72, 0.80] | 0.69 [0.59, 0.76] | 0.42 [0.19, 0.62] |
| ThermoMPNN stability (−ΔΔ*G*) | 0.41 [0.31, 0.50] | 0.63 [0.54, 0.70] | 0.65 [0.54, 0.72] | 0.70 [0.55, 0.80] |
| RSA | 0.41 [0.26, 0.53] | 0.59 [0.49, 0.65] | 0.56 [0.42, 0.64] | 0.52 [0.34, 0.66] |
| base model (ESM2+RSA+nbr) | 0.76 [0.69, 0.80] | 0.81 [0.77, 0.84] | 0.76 [0.69, 0.81] | 0.69 [0.57, 0.80] |
| full model (ESM2+RSA+nbr+ΔΔ*G*) | 0.78 [0.71, 0.82] | 0.86 [0.82, 0.87] | 0.82 [0.77, 0.85] | 0.79 [0.70, 0.86] |
| gain from adding ΔΔ*G* (full − base) | 0.02 [0.00, 0.03] | 0.04 [0.02, 0.07] | 0.06 [0.03, 0.09] | 0.10 [0.04, 0.20] |
| *TEM-1 Firnberg 2014* |  |  |  |  |
| ESM2 LLR | 0.74 [0.66, 0.79] | 0.77 [0.72, 0.80] | 0.70 [0.60, 0.77] | 0.40 [0.16, 0.60] |
| ThermoMPNN stability (−ΔΔ*G*) | 0.44 [0.34, 0.54] | 0.61 [0.53, 0.69] | 0.64 [0.53, 0.73] | 0.72 [0.56, 0.80] |
| RSA | 0.44 [0.30, 0.56] | 0.62 [0.53, 0.68] | 0.56 [0.42, 0.64] | 0.49 [0.28, 0.65] |
| base model (ESM2+RSA+nbr) | 0.80 [0.75, 0.84] | 0.83 [0.79, 0.85] | 0.76 [0.69, 0.82] | 0.67 [0.53, 0.78] |
| full model (ESM2+RSA+nbr+ΔΔ*G*) | 0.82 [0.77, 0.86] | 0.86 [0.82, 0.88] | 0.81 [0.76, 0.85] | 0.78 [0.66, 0.85] |
| gain from adding ΔΔ*G* (full − base) | 0.02 [0.00, 0.03] | 0.03 [0.02, 0.05] | 0.05 [0.02, 0.08] | 0.11 [0.03, 0.20] |
| *TEM-1 Jacquier 2013* |  |  |  |  |
| ESM2 LLR | 0.79 [0.73, 0.84] | 0.72 [0.66, 0.78] | 0.61 [0.47, 0.72] | 0.32 [0.04, 0.57] |
| ThermoMPNN stability (−ΔΔ*G*) | 0.45 [0.33, 0.57] | 0.54 [0.42, 0.63] | 0.53 [0.39, 0.64] | 0.57 [0.38, 0.70] |
| RSA | 0.31 [0.13, 0.47] | 0.48 [0.36, 0.58] | 0.38 [0.23, 0.49] | 0.47 [0.30, 0.61] |
| base model (ESM2+RSA+nbr) | 0.82 [0.75, 0.86] | 0.75 [0.69, 0.80] | 0.63 [0.49, 0.72] | 0.61 [0.45, 0.72] |
| full model (ESM2+RSA+nbr+ΔΔ*G*) | 0.82 [0.76, 0.86] | 0.77 [0.71, 0.81] | 0.65 [0.53, 0.74] | 0.62 [0.44, 0.74] |
| gain from adding ΔΔ*G* (full − base) | 0.00 [−0.01, 0.02] | 0.01 [−0.01, 0.04] | 0.02 [−0.00, 0.05] | 0.01 [−0.07, 0.09] |
| *TEM-1 Deng 2012* |  |  |  |  |
| ESM2 LLR | 0.39 [0.28, 0.50] | 0.58 [0.47, 0.67] | 0.60 [0.48, 0.67] | 0.22 [0.00, 0.43] |
| ThermoMPNN stability (−ΔΔ*G*) | 0.15 [0.01, 0.29] | 0.50 [0.40, 0.58] | 0.40 [0.27, 0.54] | 0.34 [0.12, 0.51] |
| RSA | 0.05 [−0.10, 0.21] | 0.37 [0.22, 0.49] | 0.32 [0.16, 0.46] | 0.19 [−0.01, 0.38] |
| base model (ESM2+RSA+nbr) | 0.33 [0.22, 0.44] | 0.58 [0.44, 0.67] | 0.62 [0.48, 0.71] | 0.33 [0.12, 0.51] |
| full model (ESM2+RSA+nbr+ΔΔ*G*) | 0.32 [0.21, 0.44] | 0.63 [0.51, 0.71] | 0.63 [0.50, 0.71] | 0.31 [0.07, 0.50] |
| gain from adding ΔΔ*G* (full − base) | −0.01 [−0.02, 0.01] | 0.05 [0.01, 0.09] | 0.01 [−0.01, 0.03] | −0.01 [−0.11, 0.07] |
| Predictor | <8 Å | 8–14 Å | 14–20 Å | >20 Å |
| *TEM-1 Stiffler 2015* |  |  |  |  |
| ESM2 LLR | 0.91 [0.88, 0.94] | 0.90 [0.87, 0.93] | 0.85 [0.80, 0.90] | 0.72 [0.57, 0.84] |
| ThermoMPNN stability (−ΔΔ*G*) | 0.72 [0.65, 0.78] | 0.82 [0.77, 0.87] | 0.85 [0.79, 0.90] | 0.90 [0.83, 0.95] |
| RSA | 0.71 [0.61, 0.78] | 0.81 [0.76, 0.86] | 0.82 [0.75, 0.87] | 0.79 [0.69, 0.87] |
| base model (ESM2+RSA+nbr) | 0.92 [0.89, 0.94] | 0.93 [0.90, 0.95] | 0.91 [0.87, 0.94] | 0.88 [0.82, 0.94] |
| full model (ESM2+RSA+nbr+ΔΔ*G*) | 0.93 [0.90, 0.95] | 0.95 [0.92, 0.96] | 0.94 [0.91, 0.96] | 0.94 [0.90, 0.97] |
| gain from adding ΔΔ*G* (full − base) | 0.01 [0.00, 0.01] | 0.02 [0.01, 0.03] | 0.03 [0.01, 0.05] | 0.06 [0.01, 0.12] |
| *TEM-1 Firnberg 2014* |  |  |  |  |
| ESM2 LLR | 0.90 [0.86, 0.93] | 0.90 [0.86, 0.92] | 0.85 [0.79, 0.90] | 0.70 [0.53, 0.82] |
| ThermoMPNN stability (−ΔΔ*G*) | 0.72 [0.65, 0.79] | 0.82 [0.76, 0.87] | 0.84 [0.78, 0.90] | 0.91 [0.86, 0.95] |
| RSA | 0.69 [0.59, 0.77] | 0.82 [0.76, 0.86] | 0.80 [0.73, 0.86] | 0.78 [0.68, 0.87] |
| base model (ESM2+RSA+nbr) | 0.91 [0.88, 0.94] | 0.93 [0.90, 0.95] | 0.89 [0.85, 0.93] | 0.88 [0.80, 0.93] |
| full model (ESM2+RSA+nbr+ΔΔ*G*) | 0.92 [0.89, 0.95] | 0.94 [0.92, 0.96] | 0.92 [0.89, 0.95] | 0.94 [0.90, 0.97] |
| gain from adding ΔΔ*G* (full − base) | 0.01 [0.00, 0.02] | 0.02 [0.01, 0.03] | 0.03 [0.01, 0.05] | 0.06 [0.03, 0.11] |
| *TEM-1 Jacquier 2013* |  |  |  |  |
| ESM2 LLR | 0.91 [0.87, 0.94] | 0.86 [0.81, 0.90] | 0.84 [0.76, 0.90] | 0.68 [0.51, 0.81] |
| ThermoMPNN stability (−ΔΔ*G*) | 0.73 [0.65, 0.81] | 0.78 [0.72, 0.83] | 0.79 [0.71, 0.86] | 0.84 [0.74, 0.91] |
| RSA | 0.62 [0.49, 0.73] | 0.74 [0.68, 0.80] | 0.72 [0.64, 0.78] | 0.75 [0.64, 0.84] |
| base model (ESM2+RSA+nbr) | 0.91 [0.87, 0.95] | 0.87 [0.83, 0.91] | 0.86 [0.80, 0.91] | 0.82 [0.73, 0.90] |
| full model (ESM2+RSA+nbr+ΔΔ*G*) | 0.92 [0.87, 0.95] | 0.88 [0.84, 0.92] | 0.87 [0.81, 0.91] | 0.85 [0.76, 0.91] |
| gain from adding ΔΔ*G* (full − base) | 0.00 [−0.00, 0.01] | 0.01 [−0.00, 0.03] | 0.01 [−0.01, 0.02] | 0.03 [−0.03, 0.08] |
| *TEM-1 Deng 2012* |  |  |  |  |
| ESM2 LLR | 0.72 [0.66, 0.79] | 0.81 [0.75, 0.86] | 0.80 [0.74, 0.85] | 0.63 [0.53, 0.74] |
| ThermoMPNN stability (−ΔΔ*G*) | 0.60 [0.52, 0.67] | 0.75 [0.70, 0.80] | 0.70 [0.64, 0.77] | 0.68 [0.57, 0.77] |
| RSA | 0.54 [0.47, 0.62] | 0.71 [0.64, 0.78] | 0.69 [0.60, 0.75] | 0.62 [0.51, 0.72] |
| base model (ESM2+RSA+nbr) | 0.70 [0.63, 0.76] | 0.81 [0.75, 0.86] | 0.82 [0.74, 0.87] | 0.69 [0.58, 0.78] |
| full model (ESM2+RSA+nbr+ΔΔ*G*) | 0.69 [0.62, 0.76] | 0.83 [0.78, 0.88] | 0.82 [0.74, 0.87] | 0.68 [0.56, 0.77] |
| gain from adding ΔΔ*G* (full − base) | −0.00 [−0.01, 0.00] | 0.02 [−0.00, 0.04] | 0.00 [−0.01, 0.01] | −0.01 [−0.06, 0.03] |

**Fig. 1.**
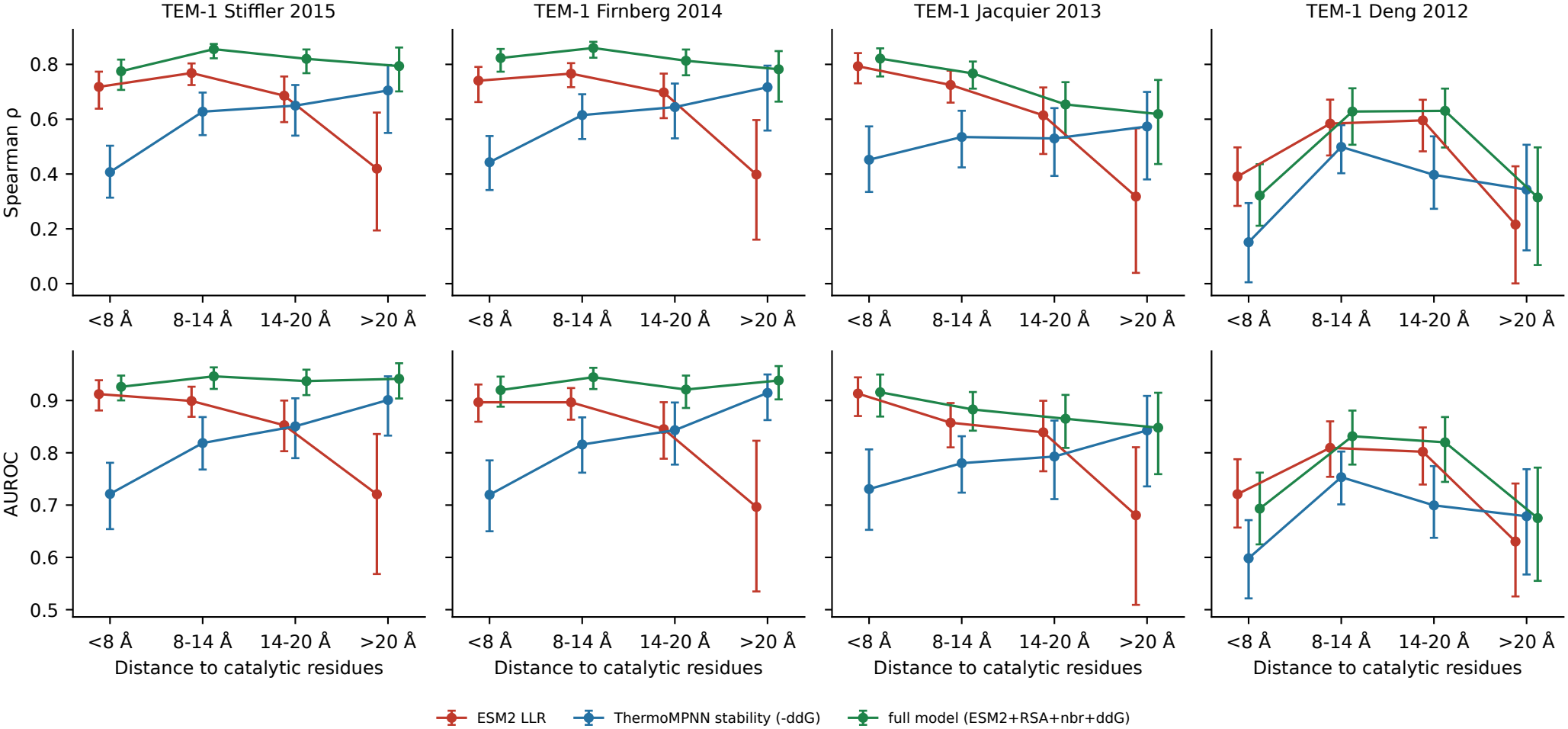
ESM2 accuracy falls and predicted stability gains information with distance in TEM-1. Spearman ρ (top) and AUROC (bottom) by distance to the catalytic residues, for four assays. ESM2 and stability are evaluated directly; the full model uses cross-validated predictions. Bars: 95% bootstrap intervals over positions

**Fig. 2.**
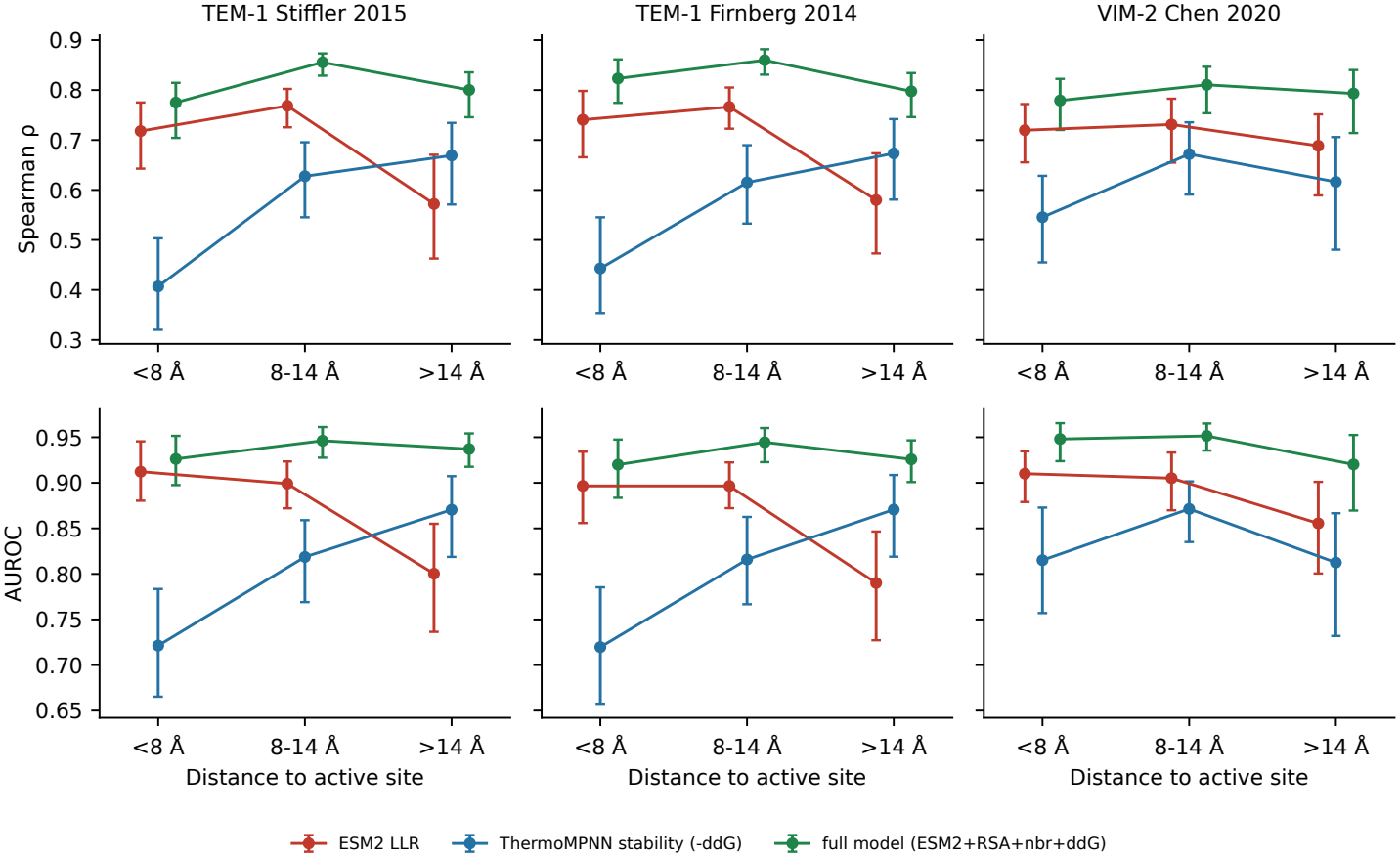
TEM-1 versus VIM-2 in matched distance bins. As Figure 1, with three bins (< 8, 8-14 and > 14 Å). VIM-2 has only seven positions beyond 20 Å, so its far bins are merged and mostly span 14-20 Å

The ThermoMPNN stability score (19) showed the opposite trend. In Stiffler its correlation with fitness rose from 0.41 below 8 Å to 0.70 (0.55–0.80) beyond 20 Å, and its AUROC from 0.72 to 0.90 (0.83–0.95; Table 2). Firnberg gave 0.44 to 0.72 and 0.72 to 0.91. The rise was smaller in Jacquier (0.45 to 0.57; AUROC 0.73 to 0.84) and unclear in Deng (0.15, 0.50, 0.40 and 0.34 across the four bins; Table S6).

Adding stability to a model of ESM2, RSA and neighbours improved cross-validated by 0.02 (0.00–0.03) below 8 Å and by 0.10 (0.04–0.20) beyond 20 Å in Stiffler. In Firnberg the gains were 0.02 and 0.11 (0.03–0.20). AUROC gains were 0.01 and 0.06 in both assays (beyond 20 Å, 95% CI 0.01– 0.12 in Stiffler and 0.03–0.11 in Firnberg). In Jacquier the gain stayed at or below 0.02, and beyond 20 Å its interval included zero (0.01, −0.07 to +0.09). The same was true in Deng (−0.01, −0.11 to +0.07; Table S6). The full model did not decline with distance in Stiffler and Firnberg: its farminus-near was +0.03 (−0.04 to +0.10) and −0.03 (−0.08 to +0.03), and its AUROC difference was +0.01 in both (Table S3). The Jacquier full model still declined (0.82 to 0.62 across the four bins; Table S6).

With three bins and with position quartiles, ESM2’s farminus-near difference was negative and the stability difference positive in Stiffler and Firnberg (Table S3). In a model with distance as a continuous variable, the ESM2 distance coefficient per standard deviation of distance (about 6 Å) was −0.142 (−0.188 to −0.094) in Stiffler, −0.138 (−0.186 to −0.094) in Firnberg and −0.170 (−0.227 to −0.116) in Jacquier. The stability distance coefficient was +0.124 (+0.076 to +0.165), +0.109 (+0.062 to +0.154) and +0.094 (+0.039 to +0.152) (Figure 3, Table S4). For Deng both intervals included zero (−0.041, −0.100 to +0.028; +0.052, −0.010 to +0.130). Removing the four catalytic residues changed none of these estimates materially (Table S4).

**Fig. 3.**
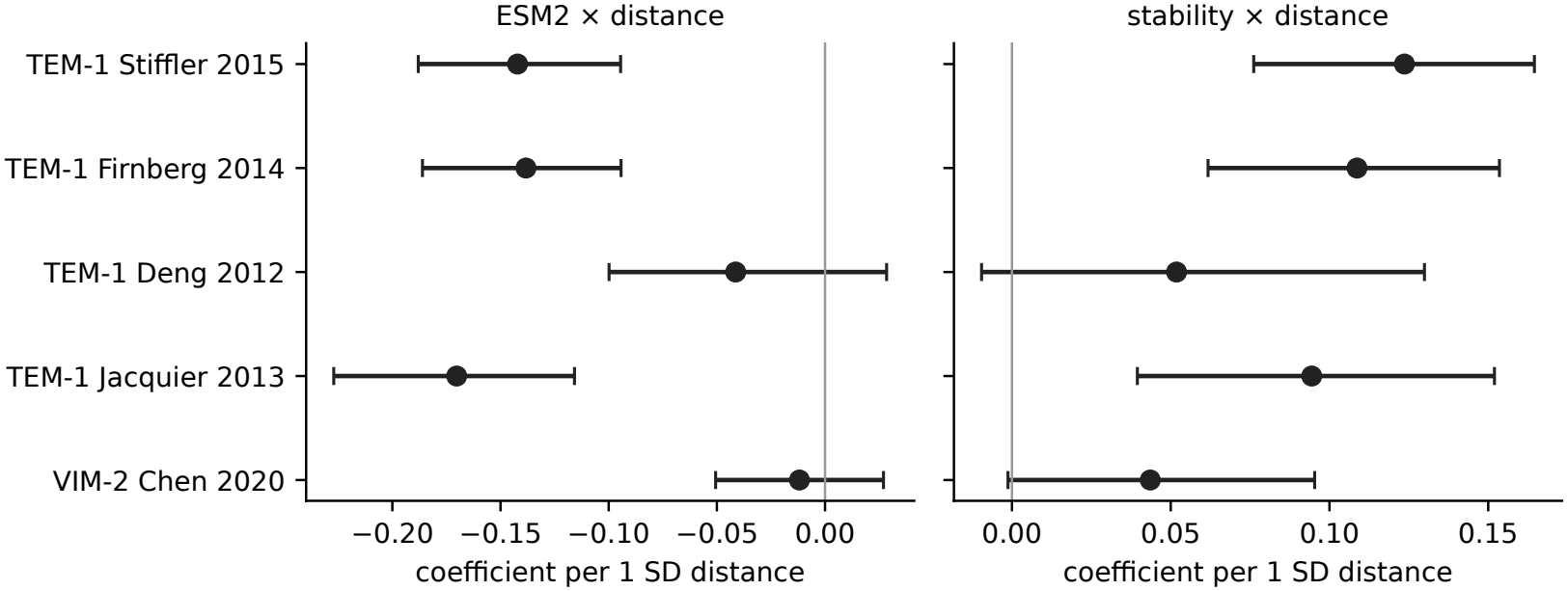
Continuous-distance model. ESM2 × distance (left) and stability × distance (right) coefficients per standard deviation of distance, with 95% bootstrap intervals. A negative ESM2 and a positive stability coefficient mean that ESM2’s weight falls and stability’s rises with distance.

The 40 TEM-1 positions beyond 20 Å lie in five contiguous stretches (Ambler 26–41, 50–59, 226–229, 251–258 and 281–290). Dropping any one stretch left ESM2’s far-minus-near difference negative (−0.22 to −0.37 in Stiffler; −0.26 to −0.41 in Firnberg) and the stability difference positive (+0.27 to +0.31; +0.24 to +0.30; Figure 4, Table S5). The ESM2 decline shrank most when 251–258 or 281–290 was removed. A block bootstrap that resamples stretches gave −0.30 (−0.62 to −0.04) for ESM2 and +0.30 (+0.19 to +0.40) for stability in Stiffler, and −0.34 (−0.63 to −0.09) and +0.27 (+0.15 to +0.41) in Firnberg (Table S5). The lower bound of the ESM2 interval in Stiffler is close to zero.

**Fig. 4.**
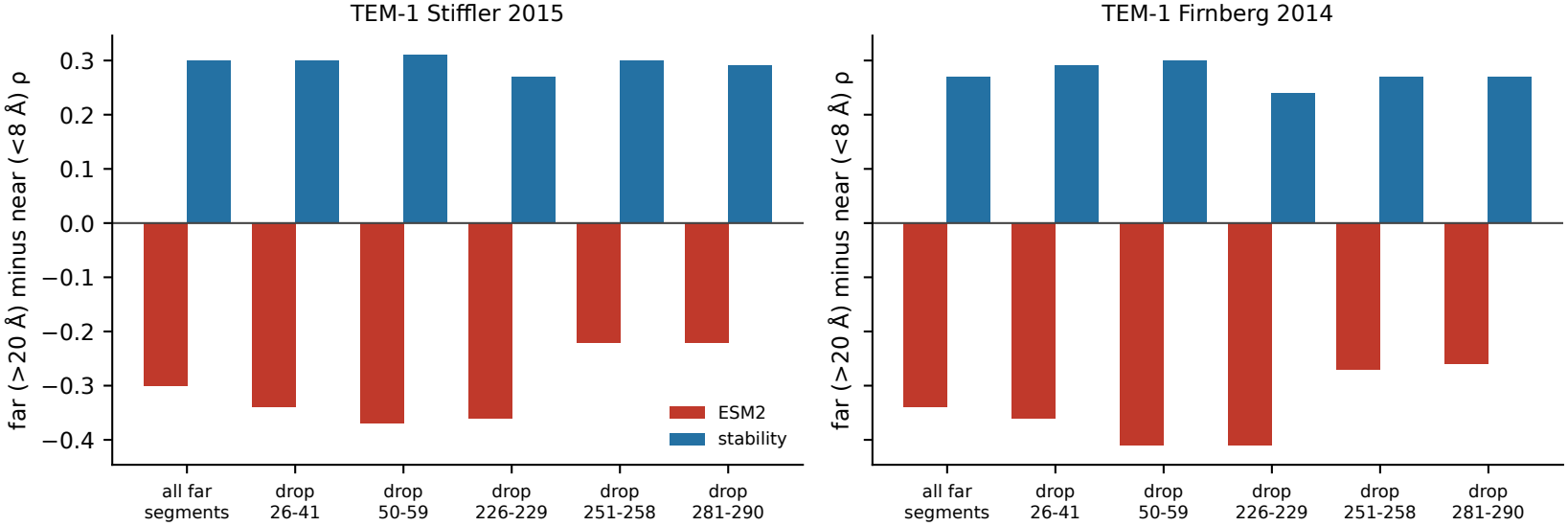
TEM-1 far-versus-near result when each stretch is dropped. Difference in Spearman ρ (> 20 Å minus < 8 Å) for ESM2 and stability, using all distal positions and dropping each of five contiguous stretches in turn. Raw correlations; block-bootstrap intervals are in Table S5

Nine positions more than 14 Å from the catalytic residues had a mean full-model residual below −0.15 in all three of the Firnberg, Stiffler and Jacquier assays: Ambler 26, 29, 197, 229, 230, 251, 259, 267 and 290 (Figure S2, Table S2). The largest were Trp290 (−0.45) and Gly251 (−0.42), at 21.6 and 20.2 Å. Their RSA ranged from 0.00 to 0.81, and five of the nine lie within the stretches beyond 20 Å. Position-level mean residuals agreed between assays (=0.86 for Firnberg versus Stiffler and 0.66–0.70 for Jacquier versus each), so these positions are unlikely to reflect noise in one assay.

VIM-2 has only seven positions beyond 20 Å from its zinc site, so we compared <8, 8–14 and >14 Å (Figure 2, Table S3). ESM2’s was 0.72, 0.73 and 0.69, with a far-minus-near difference of −0.03 (−0.13 to +0.07; AUROC −0.05, −0.11 to 0.00). The stability score’s was 0.55, 0.67 and 0.62 (difference +0.07, −0.09 to +0.20; AUROC 0.00, −0.11 to +0.08). Adding stability to the base model improved by 0.01 (−0.00 to +0.03), 0.04 (0.01–0.08) and 0.03 (0.00–0.07) in the three bins. Quartiles gave the same picture (ESM2 −0.02, −0.13 to +0.09; stability +0.09, −0.05 to +0.25; Table S3). In the continuous model the ESM2 distance coefficient was −0.012 (−0.051 to +0.027) and the stability distance coefficient +0.044 (−0.001 to +0.095). Removing the six zinc ligands left both intervals including zero (+0.012, −0.027 to +0.051; +0.033, −0.012 to +0.082; Table S4). The VIM-2 >14 Å bin resembled the TEM-1 14–20 Å bin: ESM2 was 0.69 in both, and stability was 0.62 in VIM-2 and 0.65 in Stiffler (Table 2). Of the 46 VIM-2 positions beyond 14 Å, 39 lie between 14 and 20 Å.

## 4 Discussion

In TEM-1, the accuracy of an evolutionary language model fell with distance from the catalytic residues. A structure-based stability score became more informative over the same range, and a model that combined them performed similarly at all distances. This held across binning schemes and in a continuous-distance model, and it survived the removal of any one contiguous stretch of distal positions. In VIM-2 we found no distance trend, but that dataset samples few positions beyond 20 Å, so it does not settle the question.

The pattern fits the view that many mutations far from the active site change the resistance phenotype by altering how much properly folded enzyme the cell has. Earlier work has argued that stability effects shape which mutations proteins tolerate (11), and studies of TEM-1 describe trade-offs between stability and activity and a stabilising second-site mutation that suppresses misfolding (31, 32). Analyses of DMS data from other proteins likewise suggest that about half of loss-of-function variants act through reduced stability, while sequence-based predictors are on average more accurate than stability predictors (14, 33). Predicted stability effects also correlate most strongly with abundance-based DMS phenotypes (21). The resistance phenotype combines activity, stability and expression. A predictor of folding energetics should therefore matter more where direct catalytic constraints are weak. Our data are compatible with this account but do not prove it.

Two cautions apply. First, the stability score is a prediction from one crystal structure, not a measurement. ThermoMPNN was trained on large-scale stability data of mostly small domains (18, 19), and TEM-1 is larger than most of them. The score may also carry structural information beyond folding energy, so “stability-related structural information” is the safer description. Second, the gain was moderate (+0.10 in and +0.06 in AUROC beyond 20 Å in Stiffler), and it became smaller after we corrected the solvent-accessibility calculation.

Language models such as ESM2 learn evolutionary constraint from sequence families (15). Constraint is strongest around the catalytic site, where the model does well. Further away, evolutionary constraint is weaker, yet the phenotype can still be sensitive to destabilising changes that conservation alone does not reveal. The ESM2 decline was concentrated partly in the stretches 251–258 and 281–290. Mutations in terminal or surface segments could also act through expression or export, which neither predictor models. Our data cannot separate these explanations. The VIM-2 study itself examined translocation as a separate requirement (9).

We detected no distance trend in VIM-2, but the evidence is limited. Only seven positions lie beyond 20 Å of the zinc site, and the VIM-2 far bin behaved like TEM-1’s 14–20 Å bin, where TEM-1 also looks intermediate. The enzymes also differ in catalytic class, size, selection conditions and number of replicates (VIM-2 has one assay), and ThermoMPNN does not model zinc. We did not test whether the two enzymes differ from each other. The result therefore neither supports nor contradicts a general pattern. Testing generality needs larger enzymes, or datasets with many positions beyond 20 Å from the catalytic site.

Nine distal positions were more sensitive than the full model predicted in all three of the Firnberg, Stiffler and Jacquier assays, and their residuals agreed across assays. Five of them lie in the far stretches. These positions are the best candidates for effects beyond what ESM2, burial and predicted stability capture, such as local interactions, dynamics or expression and export effects. Computational studies of TEM-1 have proposed that distal mutations can affect activity through dynamic coupling to the active site (12, 13). We did not examine their mechanisms. Targeted experiments or molecular simulations could do so.

Firnberg and Stiffler agree closely (=0.94) and are not independent confirmations of each other. The independent support rests largely on Jacquier, where the stability gain beyond 20 Å was close to zero and the full model still declined. The TEM-1 far bin contains 40 positions in five stretches, and the block-bootstrap interval for ESM2 in Stiffler approaches zero. Confidence intervals resample positions without refitting models, no multiple-comparison correction was applied, and several analysis decisions were made after seeing data (Methods), so the study is exploratory. We used one structure per enzyme (the VIM-2 structure is inhibitor-bound, with the ligand removed) and one phenotype per assay. ESM2 was trained on UniRef (34), which contains TEM-1 homologs [9], and we have not verified the overlap between ThermoMPNN’s training data and TEM-1 or VIM-2 (18, 19). Agreement between assays is only a rough indicator of reproducibility, and the Deng assay is too noisy to test the hypothesis.

## 5 Conclusions

In TEM-1, a protein language model was most accurate near the catalytic residues, and predicted stability added the most information far from them. VIM-2 did not sample the far regime well enough to test whether the pattern generalises. Larger enzymes and direct stability measurements are the next steps.

## 7 Author Contributions

Akshaya Dhakal: Conceptualization, Methodology, Investigation, Formal analysis, Visualization, and Writing – original draft. Subash Acharya: Methodology, Validation and Writing – review and editing. Gungun Chand: Literature review, Methodology, Data curation, and Writing – review and editing.

## Supplementary Material

**Table S1.** Agreement between replicate assays by distance to the catalytic residues. Fir = Firnberg, Sti = Stiffler, Jac = Jacquier, Den = Deng.

| Distance bin | Fir–Sti | Jac–Sti | Den–Sti | Den–Fir |
|---|---|---|---|---|
| <8 Å | 0.89 | 0.90 | 0.34 | 0.32 |
| 8–14 Å | 0.94 | 0.86 | 0.66 | 0.66 |
| 14–20 Å | 0.91 | 0.77 | 0.68 | 0.68 |
| >20 Å | 0.88 | 0.72 | 0.54 | 0.52 |

**Table S2.** Distal positions with consistently negative full-model residuals. Positions more than 14 Å from the catalytic residues whose mean residual was below *−*0.15 in all three assays (Firnberg, Stiffler, Jacquier). Positions are in Ambler numbering. Distance is the minimum heavy-atom distance to the catalytic residues; RSA is relative solvent accessibility; *n*_neg_ is the number of assays (out of three) in which the position’s residual was negative.

| Position | WT | Distance (Å) | RSA | Mean residual | *n*<sub>neg</sub> |
|---|---|---|---|---|---|
| 290 | W | 21.55 | 0.33 | −0.45 | 3 |
| 251 | G | 20.22 | 0.00 | −0.42 | 3 |
| 230 | F | 16.17 | 0.12 | −0.24 | 3 |
| 259 | R | 17.35 | 0.14 | −0.21 | 3 |
| 26 | H | 26.38 | 0.42 | −0.20 | 3 |
| 229 | W | 20.86 | 0.22 | −0.19 | 3 |
| 29 | T | 22.29 | 0.03 | −0.18 | 3 |
| 267 | G | 14.43 | 0.49 | −0.17 | 3 |
| 197 | E | 17.93 | 0.81 | −0.16 | 3 |

**Table S3.**
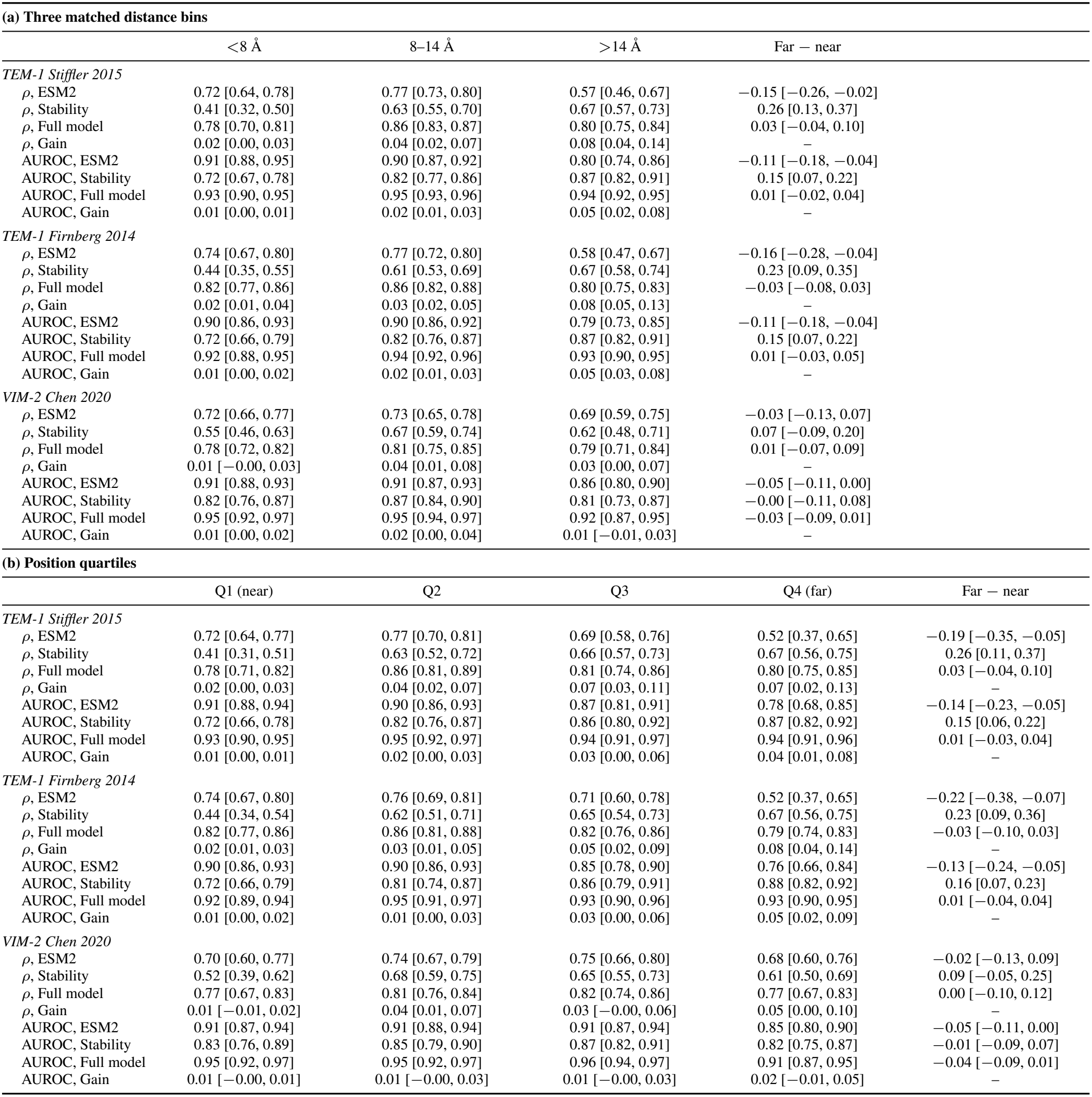
Matched-bin and quartile analyses of predictor performance by distance. Values are point estimates with 95% confidence intervals in brackets. **(a)** Three matched distance bins and **(b)** position quartiles: Spearman *ρ* and AUROC for ESM2, the stability score, and the full model, with the gain from adding stability (full *−* base) and the far-minus-near difference (last column in each panel).

| **(a) Three matched distance bins** |  |  |  |  |  |
|  | <8 Å | 8–14 Å | >14 Å | Far − near |  |
|---|---|---|---|---|---|
| *TEM-1 Stiffler 2015* |  |  |  |  |  |
| ρ, ESM2 | 0.72 [0.64, 0.78] | 0.77 [0.73, 0.80] | 0.57 [0.46, 0.67] | −0.15 [−0.26, −0.02] |  |
| ρ, Stability | 0.41 [0.32, 0.50] | 0.63 [0.55, 0.70] | 0.67 [0.57, 0.73] | 0.26 [0.13, 0.37] |  |
| ρ, Full model | 0.78 [0.70, 0.81] | 0.86 [0.83, 0.87] | 0.80 [0.75, 0.84] | 0.03 [−0.04, 0.10] |  |
| ρ, Gain | 0.02 [0.00, 0.03] | 0.04 [0.02, 0.07] | 0.08 [0.04, 0.14] | – |  |
| AUROC, ESM2 | 0.91 [0.88, 0.95] | 0.90 [0.87, 0.92] | 0.80 [0.74, 0.86] | −0.11 [−0.18, −0.04] |  |
| AUROC, Stability | 0.72 [0.67, 0.78] | 0.82 [0.77, 0.86] | 0.87 [0.82, 0.91] | 0.15 [0.07, 0.22] |  |
| AUROC, Full model | 0.93 [0.90, 0.95] | 0.95 [0.93, 0.96] | 0.94 [0.92, 0.95] | 0.01 [−0.02, 0.04] |  |
| AUROC, Gain | 0.01 [0.00, 0.01] | 0.02 [0.01, 0.03] | 0.05 [0.02, 0.08] | – |  |
| *TEM-1 Firnberg 2014* |  |  |  |  |  |
| ρ, ESM2 | 0.74 [0.67, 0.80] | 0.77 [0.72, 0.80] | 0.58 [0.47, 0.67] | −0.16 [−0.28, −0.04] |  |
| ρ, Stability | 0.44 [0.35, 0.55] | 0.61 [0.53, 0.69] | 0.67 [0.58, 0.74] | 0.23 [0.09, 0.35] |  |
| ρ, Full model | 0.82 [0.77, 0.86] | 0.86 [0.82, 0.88] | 0.80 [0.75, 0.83] | −0.03 [−0.08, 0.03] |  |
| ρ, Gain | 0.02 [0.01, 0.04] | 0.03 [0.02, 0.05] | 0.08 [0.05, 0.13] | – |  |
| AUROC, ESM2 | 0.90 [0.86, 0.93] | 0.90 [0.86, 0.92] | 0.79 [0.73, 0.85] | −0.11 [−0.18, −0.04] |  |
| AUROC, Stability | 0.72 [0.66, 0.79] | 0.82 [0.76, 0.87] | 0.87 [0.82, 0.91] | 0.15 [0.07, 0.22] |  |
| AUROC, Full model | 0.92 [0.88, 0.95] | 0.94 [0.92, 0.96] | 0.93 [0.90, 0.95] | 0.01 [−0.03, 0.05] |  |
| AUROC, Gain | 0.01 [0.00, 0.02] | 0.02 [0.01, 0.03] | 0.05 [0.03, 0.08] | – |  |
| *VIM-2 Chen 2020* |  |  |  |  |  |
| ρ, ESM2 | 0.72 [0.66, 0.77] | 0.73 [0.65, 0.78] | 0.69 [0.59, 0.75] | −0.03 [−0.13, 0.07] |  |
| ρ, Stability | 0.55 [0.46, 0.63] | 0.67 [0.59, 0.74] | 0.62 [0.48, 0.71] | 0.07 [−0.09, 0.20] |  |
| ρ, Full model | 0.78 [0.72, 0.82] | 0.81 [0.75, 0.85] | 0.79 [0.71, 0.84] | 0.01 [−0.07, 0.09] |  |
| ρ, Gain | 0.01 [−0.00, 0.03] | 0.04 [0.01, 0.08] | 0.03 [0.00, 0.07] | – |  |
| AUROC, ESM2 | 0.91 [0.88, 0.93] | 0.91 [0.87, 0.93] | 0.86 [0.80, 0.90] | −0.05 [−0.11, 0.00] |  |
| AUROC, Stability | 0.82 [0.76, 0.87] | 0.87 [0.84, 0.90] | 0.81 [0.73, 0.87] | −0.00 [−0.11, 0.08] |  |
| AUROC, Full model | 0.95 [0.92, 0.97] | 0.95 [0.94, 0.97] | 0.92 [0.87, 0.95] | −0.03 [−0.09, 0.01] |  |
| AUROC, Gain | 0.01 [0.00, 0.02] | 0.02 [0.00, 0.04] | 0.01 [−0.01, 0.03] | – |  |
| **(b) Position quartiles** |  |  |  |  |  |
|  | Q1 (near) | Q2 | Q3 | Q4 (far) | Far − near |
| *TEM-1 Stiffler 2015* |  |  |  |  |  |
| ρ, ESM2 | 0.72 [0.64, 0.77] | 0.77 [0.70, 0.81] | 0.69 [0.58, 0.76] | 0.52 [0.37, 0.65] | −0.19 [−0.35, −0.05] |
| ρ, Stability | 0.41 [0.31, 0.51] | 0.63 [0.52, 0.72] | 0.66 [0.57, 0.73] | 0.67 [0.56, 0.75] | 0.26 [0.11, 0.37] |
| ρ, Full model | 0.78 [0.71, 0.82] | 0.86 [0.81, 0.89] | 0.81 [0.74, 0.86] | 0.80 [0.75, 0.85] | 0.03 [−0.04, 0.10] |
| ρ, Gain | 0.02 [0.00, 0.03] | 0.04 [0.02, 0.07] | 0.07 [0.03, 0.11] | 0.07 [0.02, 0.13] | – |
| AUROC, ESM2 | 0.91 [0.88, 0.94] | 0.90 [0.86, 0.93] | 0.87 [0.81, 0.91] | 0.78 [0.68, 0.85] | −0.14 [−0.23, −0.05] |
| AUROC, Stability | 0.72 [0.66, 0.78] | 0.82 [0.76, 0.87] | 0.86 [0.80, 0.92] | 0.87 [0.82, 0.92] | 0.15 [0.06, 0.22] |
| AUROC, Full model | 0.93 [0.90, 0.95] | 0.95 [0.92, 0.97] | 0.94 [0.91, 0.97] | 0.94 [0.91, 0.96] | 0.01 [−0.03, 0.04] |
| AUROC, Gain | 0.01 [0.00, 0.01] | 0.02 [0.00, 0.03] | 0.03 [0.00, 0.06] | 0.04 [0.01, 0.08] | – |
| *TEM-1 Firnberg 2014* |  |  |  |  |  |
| ρ, ESM2 | 0.74 [0.67, 0.80] | 0.76 [0.69, 0.81] | 0.71 [0.60, 0.78] | 0.52 [0.37, 0.65] | −0.22 [−0.38, −0.07] |
| ρ, Stability | 0.44 [0.34, 0.54] | 0.62 [0.51, 0.71] | 0.65 [0.54, 0.73] | 0.67 [0.56, 0.75] | 0.23 [0.09, 0.36] |
| ρ, Full model | 0.82 [0.77, 0.86] | 0.86 [0.81, 0.88] | 0.82 [0.76, 0.86] | 0.79 [0.74, 0.83] | −0.03 [−0.10, 0.03] |
| ρ, Gain | 0.02 [0.01, 0.03] | 0.03 [0.01, 0.05] | 0.05 [0.02, 0.09] | 0.08 [0.04, 0.14] | – |
| AUROC, ESM2 | 0.90 [0.86, 0.93] | 0.90 [0.86, 0.93] | 0.85 [0.78, 0.90] | 0.76 [0.66, 0.84] | −0.13 [−0.24, −0.05] |
| AUROC, Stability | 0.72 [0.66, 0.79] | 0.81 [0.74, 0.87] | 0.86 [0.79, 0.91] | 0.88 [0.82, 0.92] | 0.16 [0.07, 0.23] |
| AUROC, Full model | 0.92 [0.89, 0.94] | 0.95 [0.91, 0.97] | 0.93 [0.90, 0.96] | 0.93 [0.90, 0.95] | 0.01 [−0.04, 0.04] |
| AUROC, Gain | 0.01 [0.00, 0.02] | 0.01 [0.00, 0.03] | 0.03 [0.00, 0.06] | 0.05 [0.02, 0.09] | – |
| *VIM-2 Chen 2020* |  |  |  |  |  |
| ρ, ESM2 | 0.70 [0.60, 0.77] | 0.74 [0.67, 0.79] | 0.75 [0.66, 0.80] | 0.68 [0.60, 0.76] | −0.02 [−0.13, 0.09] |
| ρ, Stability | 0.52 [0.39, 0.62] | 0.68 [0.59, 0.75] | 0.65 [0.55, 0.73] | 0.61 [0.50, 0.69] | 0.09 [−0.05, 0.25] |
| ρ, Full model | 0.77 [0.67, 0.83] | 0.81 [0.76, 0.84] | 0.82 [0.74, 0.86] | 0.77 [0.67, 0.83] | 0.00 [−0.10, 0.12] |
| ρ, Gain | 0.01 [−0.01, 0.02] | 0.04 [0.01, 0.07] | 0.03 [−0.00, 0.06] | 0.05 [0.00, 0.10] | – |
| AUROC, ESM2 | 0.91 [0.87, 0.94] | 0.91 [0.88, 0.94] | 0.91 [0.87, 0.94] | 0.85 [0.80, 0.90] | −0.05 [−0.11, 0.00] |
| AUROC, Stability | 0.83 [0.76, 0.89] | 0.85 [0.79, 0.90] | 0.87 [0.82, 0.91] | 0.82 [0.75, 0.87] | −0.01 [−0.09, 0.07] |
| AUROC, Full model | 0.95 [0.92, 0.97] | 0.95 [0.92, 0.97] | 0.96 [0.94, 0.97] | 0.91 [0.87, 0.95] | −0.04 [−0.09, 0.01] |
| AUROC, Gain | 0.01 [−0.00, 0.01] | 0.01 [−0.00, 0.03] | 0.01 [−0.00, 0.03] | 0.02 [−0.01, 0.05] | – |

**Table S4.** Continuous-distance model coefficients. Per standard deviation of distance, with 95% intervals, for all positions and with the TEM-1 catalytic residues or VIM-2 zinc ligands removed

|  | ESM2 | Stability | ESM2 × distance | Stability × distance |
|---|---|---|---|---|
| *TEM-1 Stiffler 2015* |  |  |  |  |
| All positions | 0.498 [0.450, 0.545] | 0.303 [0.261, 0.350] | −0.142 [−0.188, −0.094] | 0.124 [0.076, 0.165] |
| Sensitivity subset | 0.493 [0.446, 0.544] | 0.307 [0.267, 0.347] | −0.131 [−0.177, −0.082] | 0.120 [0.078, 0.164] |
| *TEM-1 Firnberg 2014* |  |  |  |  |
| All positions | 0.506 [0.465, 0.553] | 0.294 [0.252, 0.337] | −0.138 [−0.186, −0.094] | 0.109 [0.062, 0.154] |
| Sensitivity subset | 0.502 [0.456, 0.556] | 0.294 [0.256, 0.332] | −0.125 [−0.167, −0.078] | 0.109 [0.061, 0.154] |
| *TEM-1 Deng 2012* |  |  |  |  |
| All positions | 0.462 [0.390, 0.539] | 0.174 [0.099, 0.247] | −0.041 [−0.100, 0.028] | 0.052 [−0.010, 0.130] |
| Sensitivity subset | 0.462 [0.397, 0.538] | 0.178 [0.099, 0.256] | −0.049 [−0.124, 0.026] | 0.047 [−0.018, 0.112] |
| *TEM-1 Jacquier 2013* |  |  |  |  |
| All positions | 0.507 [0.446, 0.572] | 0.210 [0.154, 0.276] | −0.170 [−0.227, −0.116] | 0.094 [0.039, 0.152] |
| Sensitivity subset | 0.502 [0.443, 0.572] | 0.211 [0.156, 0.274] | −0.165 [−0.217, −0.103] | 0.096 [0.044, 0.144] |
| *VIM-2 Chen 2020* |  |  |  |  |
| All positions | 0.505 [0.465, 0.548] | 0.250 [0.192, 0.306] | −0.012 [−0.051, 0.027] | 0.044 [−0.001, 0.095] |
| Sensitivity subset | 0.493 [0.449, 0.536] | 0.257 [0.200, 0.315] | 0.012 [−0.027, 0.051] | 0.033 [−0.012, 0.082] |

**Table S5.** TEM-1 far-versus-near differences when each stretch is dropped. Difference in raw Spearman ρ (> 20 Å minus < 8 Å) for ESM2 and stability, with block-bootstrap intervals over stretches

| Analysis | ESM2 | Stability |
|---|---|---|
| *TEM-1 Stiffler 2015* |  |  |
| all far segments (raw *ρ* difference) | −0.30 | 0.30 |
| drop segment 26–41 | −0.34 | 0.30 |
| drop segment 50–59 | −0.37 | 0.31 |
| drop segment 226–229 | −0.36 | 0.27 |
| drop segment 251–258 | −0.22 | 0.30 |
| drop segment 281–290 | −0.22 | 0.29 |
| block bootstrap over segments (95% CI) | −0.30 [−0.62, −0.04] | 0.30 [0.19, 0.40] |
| *TEM-1 Firnberg 2014* |  |  |
| all far segments (raw *ρ* difference) | −0.34 | 0.27 |
| drop segment 26–41 | −0.36 | 0.29 |
| drop segment 50–59 | −0.41 | 0.30 |
| drop segment 226–229 | −0.41 | 0.24 |
| drop segment 251–258 | −0.27 | 0.27 |
| drop segment 281–290 | −0.26 | 0.27 |
| block bootstrap over segments (95% CI) | −0.34 [−0.63, −0.09] | 0.27 [0.15, 0.41] |

**Table S6.** Performance of each predictor by distance to the catalytic residues for the TEM-1 Jacquier 2013 and Deng 2012 assays: Spearman *ρ* and AUROC with 95% confidence intervals in brackets.

| Predictor | <8 Å | 8–14 Å | 14–20 Å | >20 Å |
|---|---|---|---|---|
| *TEM-1 Jacquier 2013* |  |  |  |  |
| Spearman *ρ* |  |  |  |  |
| ESM2 LLR | 0.79 [0.73, 0.84] | 0.72 [0.66, 0.78] | 0.61 [0.47, 0.72] | 0.32 [0.04, 0.57] |
| ThermoMPNN stability (−ΔΔ*G*) | 0.45 [0.33, 0.57] | 0.54 [0.42, 0.63] | 0.53 [0.39, 0.64] | 0.57 [0.38, 0.70] |
| RSA | 0.31 [0.13, 0.47] | 0.48 [0.36, 0.58] | 0.38 [0.23, 0.49] | 0.47 [0.30, 0.61] |
| Base model (ESM2+RSA+nbr) | 0.82 [0.75, 0.86] | 0.75 [0.69, 0.80] | 0.63 [0.49, 0.72] | 0.61 [0.45, 0.72] |
| Full model (ESM2+RSA+nbr+ΔΔ*G*) | 0.82 [0.76, 0.86] | 0.77 [0.71, 0.81] | 0.65 [0.53, 0.74] | 0.62 [0.44, 0.74] |
| Gain from adding ΔΔ*G* (full − base) | 0.00 [−0.01, 0.02] | 0.01 [−0.01, 0.04] | 0.02 [−0.00, 0.05] | 0.01 [−0.07, 0.09] |
| AUROC |  |  |  |  |
| ESM2 LLR | 0.91 [0.87, 0.94] | 0.86 [0.81, 0.90] | 0.84 [0.76, 0.90] | 0.68 [0.51, 0.81] |
| ThermoMPNN stability (−ΔΔ*G*) | 0.73 [0.65, 0.81] | 0.78 [0.72, 0.83] | 0.79 [0.71, 0.86] | 0.84 [0.74, 0.91] |
| RSA | 0.62 [0.49, 0.73] | 0.74 [0.68, 0.80] | 0.72 [0.64, 0.78] | 0.75 [0.64, 0.84] |
| Base model (ESM2+RSA+nbr) | 0.91 [0.87, 0.95] | 0.87 [0.83, 0.91] | 0.86 [0.80, 0.91] | 0.82 [0.73, 0.90] |
| Full model (ESM2+RSA+nbr+ΔΔ*G*) | 0.92 [0.87, 0.95] | 0.88 [0.84, 0.92] | 0.87 [0.81, 0.91] | 0.85 [0.76, 0.91] |
| Gain from adding ΔΔ*G* (full − base) | 0.00 [−0.00, 0.01] | 0.01 [−0.00, 0.03] | 0.01 [−0.01, 0.02] | 0.03 [−0.03, 0.08] |
| *TEM-1 Deng 2012* |  |  |  |  |
| Spearman *ρ* |  |  |  |  |
| ESM2 LLR | 0.39 [0.28, 0.50] | 0.58 [0.47, 0.67] | 0.60 [0.48, 0.67] | 0.22 [0.00, 0.43] |
| ThermoMPNN stability (−ΔΔ*G*) | 0.15 [0.01, 0.29] | 0.50 [0.40, 0.58] | 0.40 [0.27, 0.54] | 0.34 [0.12, 0.51] |
| RSA | 0.05 [−0.10, 0.21] | 0.37 [0.22, 0.49] | 0.32 [0.16, 0.46] | 0.19 [−0.01, 0.38] |
| Base model (ESM2+RSA+nbr) | 0.33 [0.22, 0.44] | 0.58 [0.44, 0.67] | 0.62 [0.48, 0.71] | 0.33 [0.12, 0.51] |
| Full model (ESM2+RSA+nbr+ΔΔ*G*) | 0.32 [0.21, 0.44] | 0.63 [0.51, 0.71] | 0.63 [0.50, 0.71] | 0.31 [0.07, 0.50] |
| Gain from adding ΔΔ*G* (full − base) | −0.01 [−0.02, 0.01] | 0.05 [0.01, 0.09] | 0.01 [−0.01, 0.03] | −0.01 [−0.11, 0.07] |
| AUROC |  |  |  |  |
| ESM2 LLR | 0.72 [0.66, 0.79] | 0.81 [0.75, 0.86] | 0.80 [0.74, 0.85] | 0.63 [0.53, 0.74] |
| ThermoMPNN stability (−ΔΔ*G*) | 0.60 [0.52, 0.67] | 0.75 [0.70, 0.80] | 0.70 [0.64, 0.77] | 0.68 [0.57, 0.77] |
| RSA | 0.54 [0.47, 0.62] | 0.71 [0.64, 0.78] | 0.69 [0.60, 0.75] | 0.62 [0.51, 0.72] |
| Base model (ESM2+RSA+nbr) | 0.70 [0.63, 0.76] | 0.81 [0.75, 0.86] | 0.82 [0.74, 0.87] | 0.69 [0.58, 0.78] |
| Full model (ESM2+RSA+nbr+ΔΔ*G*) | 0.69 [0.62, 0.76] | 0.83 [0.78, 0.88] | 0.82 [0.74, 0.87] | 0.68 [0.56, 0.77] |
| Gain from adding ΔΔ*G* (full − base) | −0.00 [−0.01, 0.00] | 0.02 [−0.00, 0.04] | 0.00 [−0.01, 0.01] | −0.01 [−0.06, 0.03] |

**Fig. S1.**
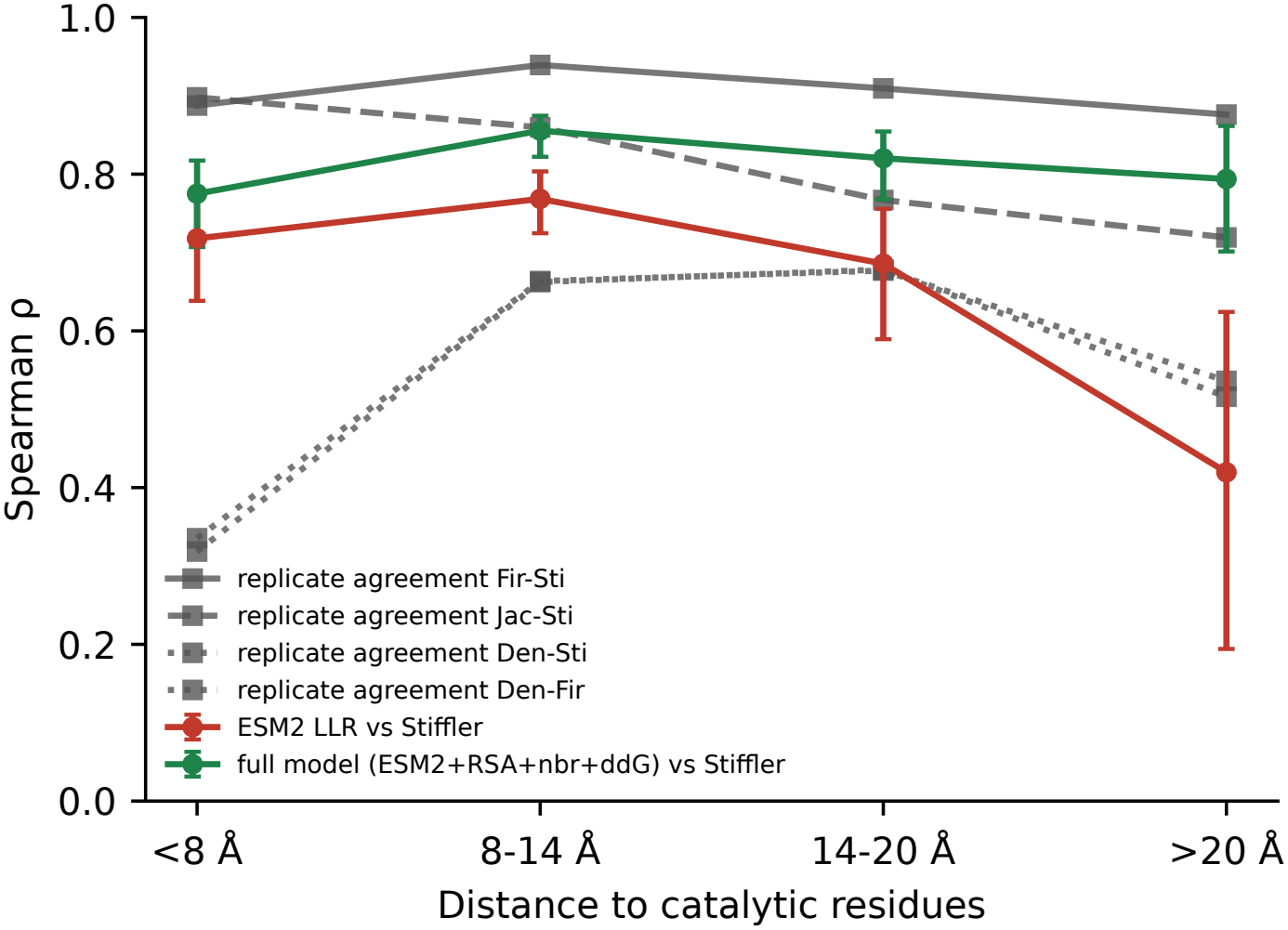
Agreement between TEM-1 assays by distance. Grey: Spearman ρ between assay pairs on shared variants. Red and green: ESM2 and the full model against Stiffler. A rough guide to reproducibility, not an exact ceiling

**Fig. S2.**
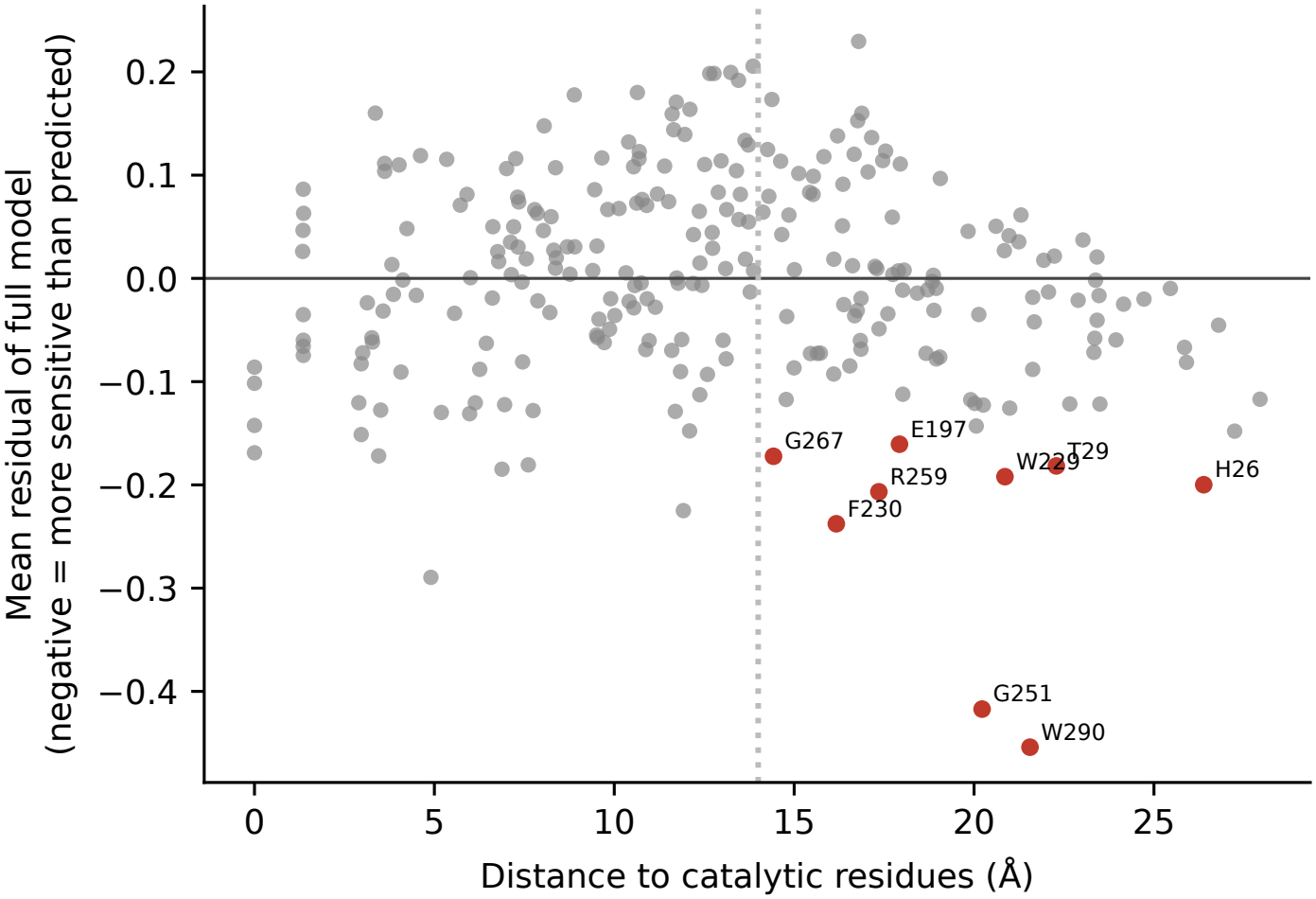
Distal positions more sensitive than predicted. Mean full-model residual (averaged over Firnberg, Stiffler and Jacquier; negative means more sensitive) against distance. Red: positions beyond 14 Å with a mean residual below −0.15 and negative in all three assays (Ambler numbers)

